# Structural identification of a selectivity filter in CFTR

**DOI:** 10.1101/2023.10.11.561906

**Authors:** Jesper Levring, Jue Chen

## Abstract

The cystic fibrosis transmembrane conductance regulator (CFTR) is a chloride channel that regulates transepithelial salt and fluid homeostasis. CFTR dysfunction leads to reduced chloride secretion into the mucosal lining of epithelial tissues, thereby causing the inherited disease cystic fibrosis. Although several structures of CFTR are available, our understanding of the ion-conduction pathway is incomplete. In particular, the route that connects the cytosolic vestibule with the extracellular space has not been clearly defined, and the structure of the open pore remains elusive. Furthermore, although many residues have been implicated in altering the selectivity of CFTR, the structure of the “selectivity filter” has yet to be determined. In this study, we identify a chloride-binding site at the extracellular ends of transmembrane helices 1, 6, and 8, where a dehydrated chloride is coordinated by residues G103, R334, F337, T338, and Y914. Alterations to this site, consistent with its function as a selectivity filter, affect ion selectivity, conductance, and open channel block. The selectivity filter is accessible from the cytosol through a large inner vestibule and opens to the extracellular solvent through a narrow portal. The identification of a chloride-binding site at the intra- and extracellular bridging point leads us to propose a complete conductance path that permits dehydrated chloride ions to traverse the lipid bilayer.

**Significance statement:** Cystic fibrosis is a fatal disease caused by inherited defects in the *cftr* gene. Understanding the structure and function of the CFTR protein is crucial for cystic fibrosis research. As an ion channel evolved from a family of ATP-driven active transporters, CFTR is structurally distinct from any other ion channel. This study describes the structure of CFTR’s ‘selectivity filter’, which enables us to complete the molecular description of the CFTR pore. Moreover, it enriches our broader knowledge of ion channel physiology, with a particular focus on chloride permeation mechanisms.

## Introduction

The cystic fibrosis transmembrane conductance regulator (CFTR), a phosphorylation- and ATP-gated anion channel (1–6), functions to regulate salt and fluid homeostasis across epithelial membranes (7). Alterations to CFTR that interfere with expression, folding, or localization to the plasma membrane or the function of the mature channel cause reduced chloride and bicarbonate transport across the apical surface of epithelial tissues and lead to the autosomal recessive disorder cystic fibrosis (7–14). CFTR hyperactivation during infection with *Vibrio cholerae* or enterotoxigenic *Escherichia coli* leads to secretory diarrhea and is central to pathogenesis in autosomal dominant polycystic kidney disease (ADPKD) (15–18). Extensive effort has been devoted to characterizing CFTR folding, gating, and ion conduction (19, 20) and to develop pharmacological modulators of these processes (21–27).

CFTR belongs to the ATP-binding cassette (ABC) transporter superfamily. Like other ABC transporters, CFTR consists of two nucleotide binding domains (NBDs) and two transmembrane domains (TMDs) (19, 20). In addition, CFTR also contains a cytosolic regulatory (R) domain bearing more than 10 predicted phosphorylation sites (28). Protein kinase A (PKA)-phosphorylation of the R domain is necessary for channel activation (2–5). Once phosphorylated, ATP-dependent dimerization of the NBDs is coupled to movements of the TMDs that open the pore (1, 6, 29, 30). Hydrolysis of ATP in CFTR’s catalytically competent ATP-binding site leads to pore closure (6, 31).

Most ABC transporters are ATP-driven pumps that function via the alternating access model, in which a translocation pathway opens only to one side of the membrane at a time (32–34). In NBD-dimerized (outward-facing) structures of typical ABC transporters, the substrate binding site is occluded from exchange with the cytosol (35–37). By contrast, in the NBD-dimerized structure of CFTR, characterized using the hydrolysis-deficient E1371Q variant, the cytosol is continuous with an inner vestibule through a lateral portal between TMs 4 and 6 through which hydrated chloride can freely diffuse (29, 38–40). This structural feature renders CFTR an ion channel instead of an active transporter, as it encloses a continuous conduit across the membrane for chloride to permeate down its electrochemical gradient.

Decades of mutagenesis studies have indicated that the pore in CFTR has an overall shape of a distorted hourglass, consisting of a shallow outer vestibule and a deep cytosolic vestibule, separated by a narrow constriction (reviewed by Linsdell (41)). Whereas the intracellular entrance (38, 39) and the cytosolic vestibule (42–46) of the pore were clearly defined in the published structures of CFTR (40, 47–49), the exact path of chloride from the inner vestibule to the extracellular space remains unclear. Extensive mapping by mutagenesis (50–53), cysteine accessibility (38, 54–56), and molecular dynamics (57) has also not allowed unambiguous assignment of the extracellular exit. Furthermore, although many residues have been shown to influence the conductance or relative permeabilities of different anions in CFTR (reviewed in (19, 20, 41)), a specific selectivity filter has yet to be defined.

In this study, we analyzed the 2.7 Å cryo-EM map of the E1371Q variant and identified a chloride-binding site at the extracellular end of the pore, where a dehydrated chloride ion is coordinated by G103, R334, F337, T338, and Y914. Electrophysiological measurements indicate that this binding site functions as a selectivity filter, determining the relative permeabilities of different anions. The selectivity filter is accessible from the cytosol and connected to the extracellular space through a narrow portal between TMs 1 and 6. These data identify the complete ion permeation path in CFTR and suggest that the structure of the NBD-dimerized E1371Q variant reflects a pore-open state rather than a flicker-closed state.

## Results

### Identification of a chloride-binding site at the extracellular tip of the pore

Previously we have reported the structure of a CFTR variant with high open probability (E1371Q) bound to the type I folding corrector lumacaftor to a resolution of 2.7 Å (48). In this structure, the two NBDs have formed a canonical ATP-dependent dimer, which is coupled to isomerization in the TMDs necessary for pore opening (Figure 1A). Mapping the dimensions of the channel using a spherical probe with a radius equal to a dehydrated chloride ion (1.7 Å) clearly identifies a solvent accessible vestibule in which ions can diffuse from the cytosol to the extracellular surface of the membrane (Figure 1A, blue surface). Close inspection of the 2.7 Å map revealed a strong density at the tip of this vestibule which likely corresponds to an ion (Figure 1B). The density is observed in an area of the map with a high degree of completeness, and it appears at a higher contour than other densities attributed to map noise.

**Figure 1.**
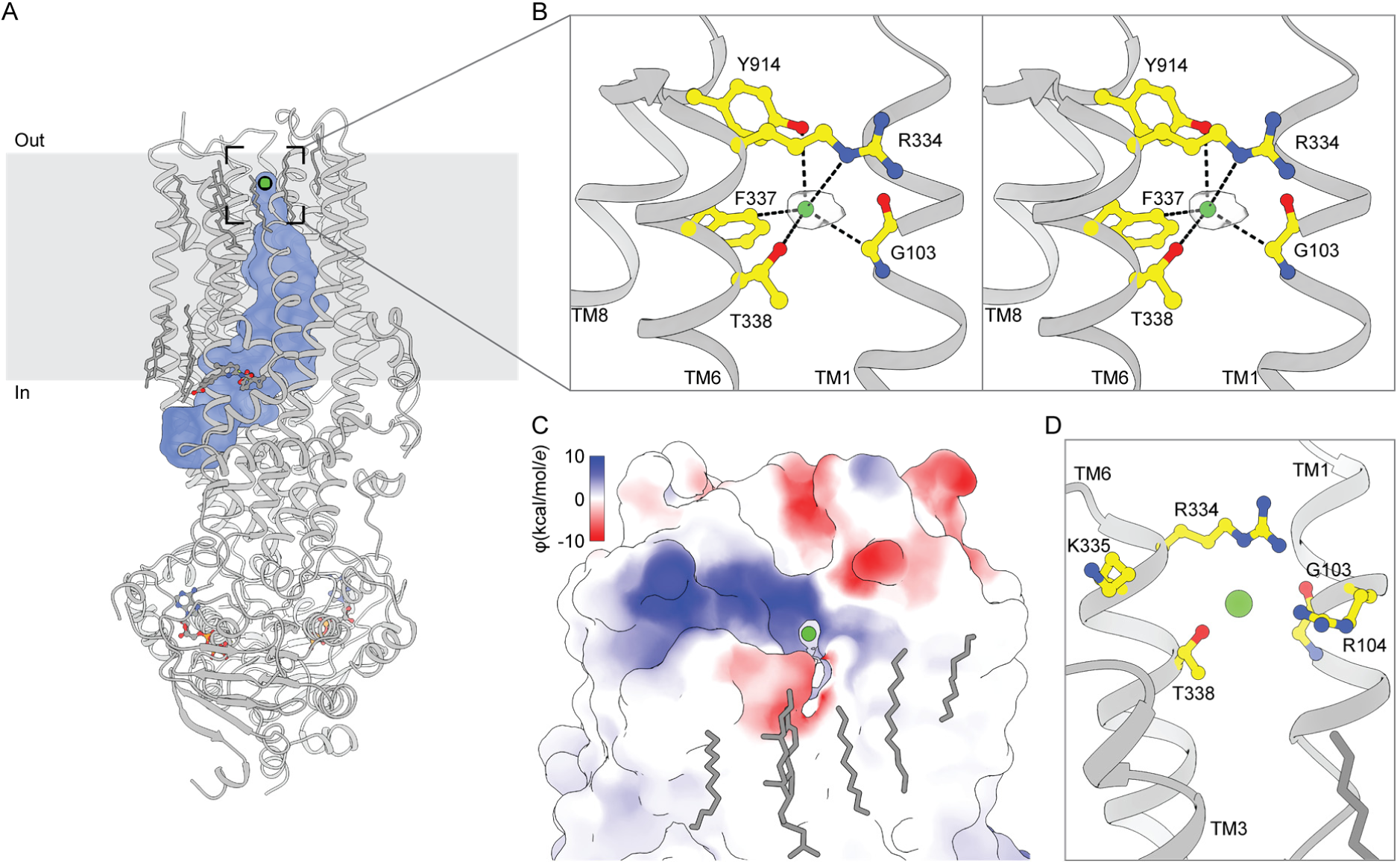
An anion selectivity filter in CFTR’s pore. (**A**) Structure of the ATP- and lumacaftor-bound E1371Q variant CFTR (Protein Data Bank 7SVD). The pore dimensions were mapped using a spherical probe with a radius equal to a dehydrated chloride ion (1.7 Å) and are shown as a blue surface. (**B**) Stereo view of chloride, shown as a green sphere within the experimental density, bound in the CFTR selectivity filter. (**C**) Surface representation of CFTR showing the extracellular exit from the selectivity filter. The surface is colored by electrostatic potential. (**D**) Closeup view from the same angle as in **C** of residues lining the extracellular exit from the selectivity filter.

The putative ion density is surrounded by residues suitable for chloride coordination (58). The hydroxyls of T338 and Y914 engage in anion–dipole interactions with coordination lengths of 3.4 Å from O^γ1^ of T338 and 3.8 Å from O^η^ of Y914 (Figure 1B). The N^ε^ of R334 is positioned 4.9 Å away to engage in a long range salt bridge that may stabilize the negative charge of permeating anions. The aromatic side chain of F337 directs its electropositive edge towards the chloride, with C^δ^ 3.4 Å away to engage in an anion–quadrupole interaction (59). In addition, the alpha carbon of G103 is 3.6 Å away, which may further stabilize the bound ion. Because the most abundant anion in the cryo-EM sample is chloride (present at 200 mM), and because the coordination geometry is consistent with chloride (58), we assigned the density to a dehydrated chloride ion.

The most prominent path from the chloride-binding site to the extracellular milieu is through a narrow lateral exit between TMs 1 and 6, delimited by the side chains of R104, R334, K335, and T338 and the main chain of G103 (Figure 1C-D). The electropositive potential around the external region of the opening is consistent with it being an anion conduction pathway, stabilizing anions through electrostatic interactions. Previously, based on a lower resolution structure obtained in the absence of lumacaftor, we proposed that this opening to be the extracellular mouth of the pore (40). The identification of a bound chloride ion at its junction with the inner vestibule lends further support for this hypothesis.

Residues that coordinate the chloride or line the extracellular mouth are highly conserved and many have been previously implicated in ion conduction or selectivity. These include the directly coordinating R334, F337, T338, and Y914 (60–65) and the residues L102, K335, and S341 located immediately proximal to the site (53, 61, 64, 66). Defective conductance has also been associated with the cystic fibrosis-causative R334W, R334L, and T338I missense mutations (14, 51, 67, 68). Combining these data with structural observations lead us to hypothesize that the observed site, termed S_filter_, corresponds to a selectivity filter that is transited by anions during conduction.

### Altered anion permeabilities of selectivity filter variants

To further test the functional importance of S_filter_, we substituted the contributing residues individually and analyzed if the properties of the pore were altered. Substitution of residues at the putative anion-binding site did not affect expression or folding of CFTR, evident from the size-exclusion chromatography profiles of all tested variants, which closely resembled that of the wild-type CFTR (Figure S1). Nonetheless, we were unable to measure currents from G103D (7 trials), G103I (10 trials), and Y914A (23 trials) variants, potentially due to complete occlusion or distortion of the chloride permeation path.

For the S_filter_ site variants from which currents were measurable, we first tested whether the site acts as a selectivity filter by measuring the effect of substitutions on relative anion permeabilities. Relative permeabilities of fluoride, chloride, bromide, iodide and thiocyanate were estimated from reversal potentials in excised inside-out patches under biionic conditions using the Goldman-Hodgkin-Katz equation (69, 70) (Figure 2 and Figure S2). In these experiments, inside-out patches were excised from Chinese Hamster Ovary (CHO) cells transiently transfected with the wild-type CFTR or S_filter_ variants. CFTR was then activated by PKA-phosphorylation, currents elicited by ATP stimulation, and current-voltage relationships measured while perfusing different permeant anions onto the patch (Figure 2A). As was previously reported (63, 71), and consistent with permeating anions having to dehydrate, wild-type CFTR exhibits a lyotropic permeability sequence, with relative permeabilities inversely related to the enthalpy of dehydration (72) (Figure 2B). Upon R334A, F337A, T338A, or Y914F substitution, the relative anion permeabilities were all altered, albeit to different degrees (Figure 2A-B). Specifically, the R334A substitution reduced the relative permeabilities for bromide and thiocyanate. The F337A substitution rendered CFTR much less discriminating against fluoride and substantially increased fluoride conductance. The T338A substitution only caused subtle changes, with an increased relative fluoride permeability being most prominent. T338A substitution also made the iodide and thiocyanate conductances more similar to chloride conductance than in the case of wild-type CFTR (Figure 2A). Most relative permeabilities of the Y914F variant were similar to the wild-type CFTR, but the relative bromide permeability was subtly reduced. These data strongly support that the ion-binding site comprising R334, F337, T338, and Y914 functions as a selectivity filter, determining the relative permeabilities of different anions.

**Figure 2.**
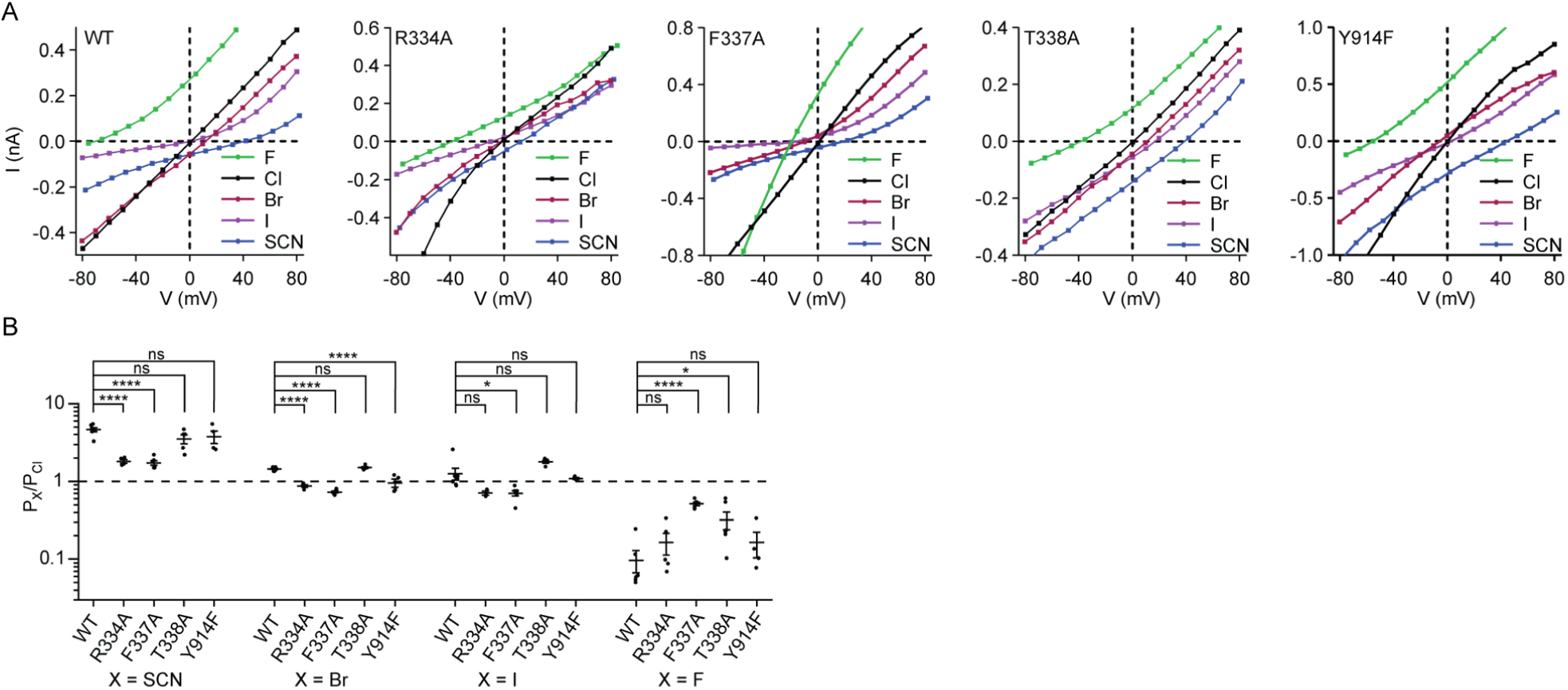
Anion permeabilities of selectivity filter variants. (**A**) Example current-voltage relationships for PKA-phosphorylated CFTR variants under biionic conditions in inside-out excised patches. The patch pipette contained 150 mM chloride, and the perfusion solution contained 150 mM of the indicated anion. 3 mM ATP was included in the perfusion solution. Membrane potentials are indicated using physiological convention. Corresponding voltage-families are in **Figure S2**. (**B**) Permeabilities of anions relative to chloride for the wild-type CFTR and selectivity filter variants. Data represent means and standard errors for 3 to 7 patches. Statistical significance was tested by one-way analysis of variance (ns: not significant, \**p* < 0.05, \*\*\*\**p* < 10^-4^).

### Reduced thiocyanate block of selectivity filter variants

Whereas thiocyanate is a permeant ion, it also binds more tightly to the CFTR pore than chloride and therefore inhibits chloride conductance (52, 62, 73). To test whether thiocyanate block occurs at S_filter_, we measured the effects of thiocyanate on the conductance of S_filter_ variants (Figure 3 and Figure S3). Specifically, we measured current-voltage relationships with different mole fractions of thiocyanate perfused onto inside-out patches excised from CHO cells expressing each of the selectivity filter variants (Figure 3A). Thiocyanate block was quantified as the relative reduction of conduction at the reversal potential (Figure 3B). As described in the literature (62), chloride currents measured from wild-type CFTR were substantially reduced even by low concentrations of thiocyanate (Figure 3A-B). Relative to the wild-type CFTR, the F337A, Y914F, R334A, and T338A variants all exhibited variable extents of reduced block by thiocyanate. The greatest effect was observed for the T338A variant, which became largely insensitive to thiocyanate block. The reduced efficacies of thiocyanate block towards each of the tested variants are consistent with thiocyanate binding at the S_filter_ site, competing with chloride with a greater affinity to thereby reduce conductance.

**Figure 3.**
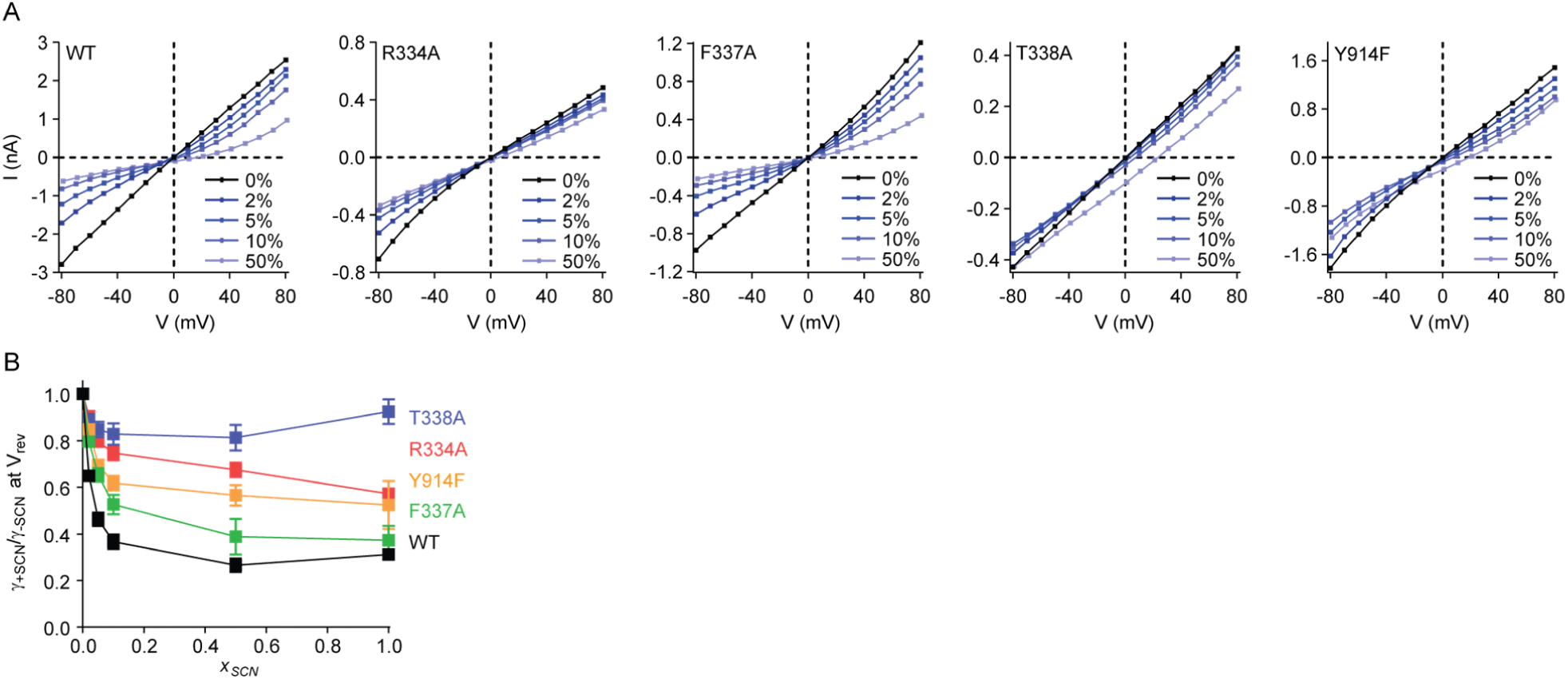
Thiocyanate block of selectivity filter variants. (**A**) Example current-voltage relationships for PKA-phosphorylated CFTR variants with different mole fractions of thiocyanate. The patch pipette contained 150 mM chloride, and the perfusion solution contained a mixture of chloride and thiocyanate. 3 mM ATP was used in the perfusion solution. Membrane potentials are indicated using physiological convention. Corresponding voltage-families are in **Figure S3**. (**B**) Thiocyanate block of CFTR conductance. Relative reduction of conduction was measured at the reversal potential as a function of the mole fraction of monovalent anion that was thiocyanate (*x_SCN_*). Data represent means and standard errors for 4-10 patches.

### Altered single-channel conductances of selectivity filter variants

We next tested the effects of substitutions at the S_filter_ site on the single-channel conductances of CFTR. Each variant was purified, phosphorylated with PKA, and reconstituted into synthetic planar lipid bilayers. The current-voltage relationships were measured in 150 mM symmetric chloride (Figure 4). Channel orientation was controlled by only including 3 mM ATP in the recording buffer of one of the two recording chambers and was validated by observing the voltage-dependence of flicker-closure events (74) (Figure 4A). Data were fitted with a parabola to account for the subtle inward rectification (Figure 4B). Under these conditions, the conductance of the wild-type CFTR at 0 mV was 7.9 ± 0.2 pS (mean and standard error). T388A substitution had little effect on single-channel conductance (8.2 ± 0.3 pS), consistent with a previous report (62). By contrast, the conductances of the three other variants were reduced to 5.7 ± 0.2 pS (Y914F), 5.0 ± 0.3 pS (F337A), and 3.0 ± 0.2 (R334A). These data indicate that disruption of interactions with the permeant ion at the S_filter_ site increase the energetic barrier for transfer through the pore, underscoring the importance of the S_filter_ for anion conduction.

**Figure 4.**
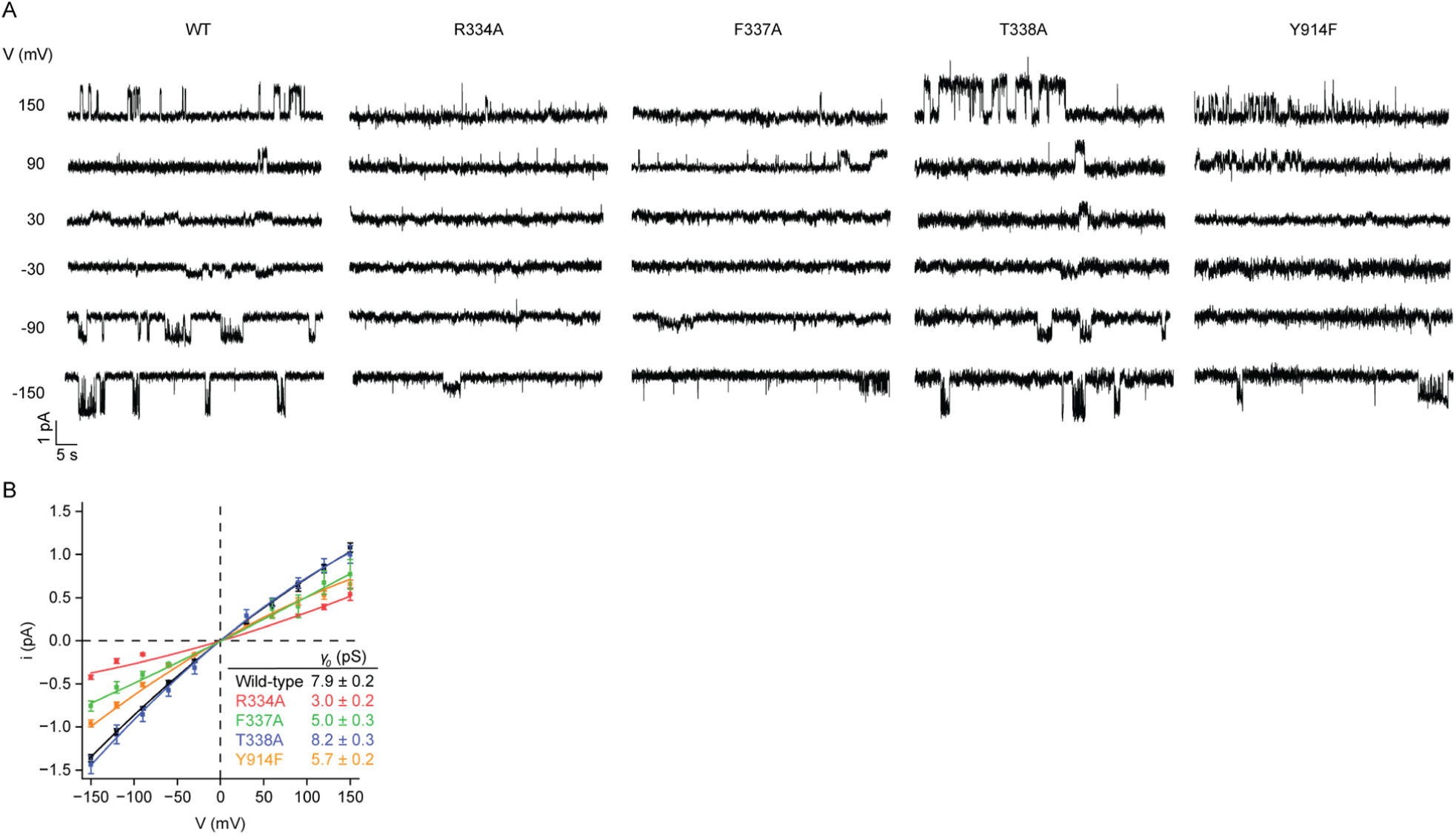
Single-channel conductances of selectivity filter variants. (**A**) Example current-voltage relationships for individual PKA-phosphorylated CFTR variants reconstituted in synthetic planar lipid bilayers. Symmetric recording buffer contained 150 mM chloride. 3 mM ATP was included in the upper recording chamber only. Membrane potentials are indicated using physiological convention. (**B**) Single-channel conductances of the wild-type CFTR and selectivity filter variants. Data represent means and standard errors for 4 to 42 channels and were fitted with quadratic functions of the form *i* = *γ*_0_*V* + *cV*^2^ to account for the subtle inward rectification. Conductances at 0 mV (*γ*_0_) are reported.

## Discussion

CFTR is an ion channel uniquely evolved from a family of active transporters. Structurally, it resembles an ABC transporter unrelated to any other ion channels. The CFTR pore has been extensively mapped with mutagenesis (reviewed in (19, 20, 41)) and partially visualized in the molecular structures of CFTR (40, 47–49). In this study, we identified a specific ion-binding site in the narrow constriction of the pore that influences ion selectivity; it also structurally connects the cytosolic vestibule with a lateral path into the extracellular space. With this information, we now have a structural understanding of the entire permeation pathway in CFTR (Figure 5A). Hydrated chloride enters the inner vestibule from the cytosol through a lateral portal between TMs 4 and 6 (Figure 5B, lower panel). Chloride remains hydrated in the inner vestibule and is stabilized by a positive electrostatic surface potential. The width of the vestibule tapers down and converges at the selectivity filter, where only dehydrated chloride can enter. Dehydrated chloride moves into the selectivity filter, stabilized by interactions with G103, R334, F337, T338, and Y914, and rehydrates upon exit into the epithelial lumen through a narrow lateral exit between TMs 1 and 6 (Figure 5B, upper panel).

**Figure 5.**
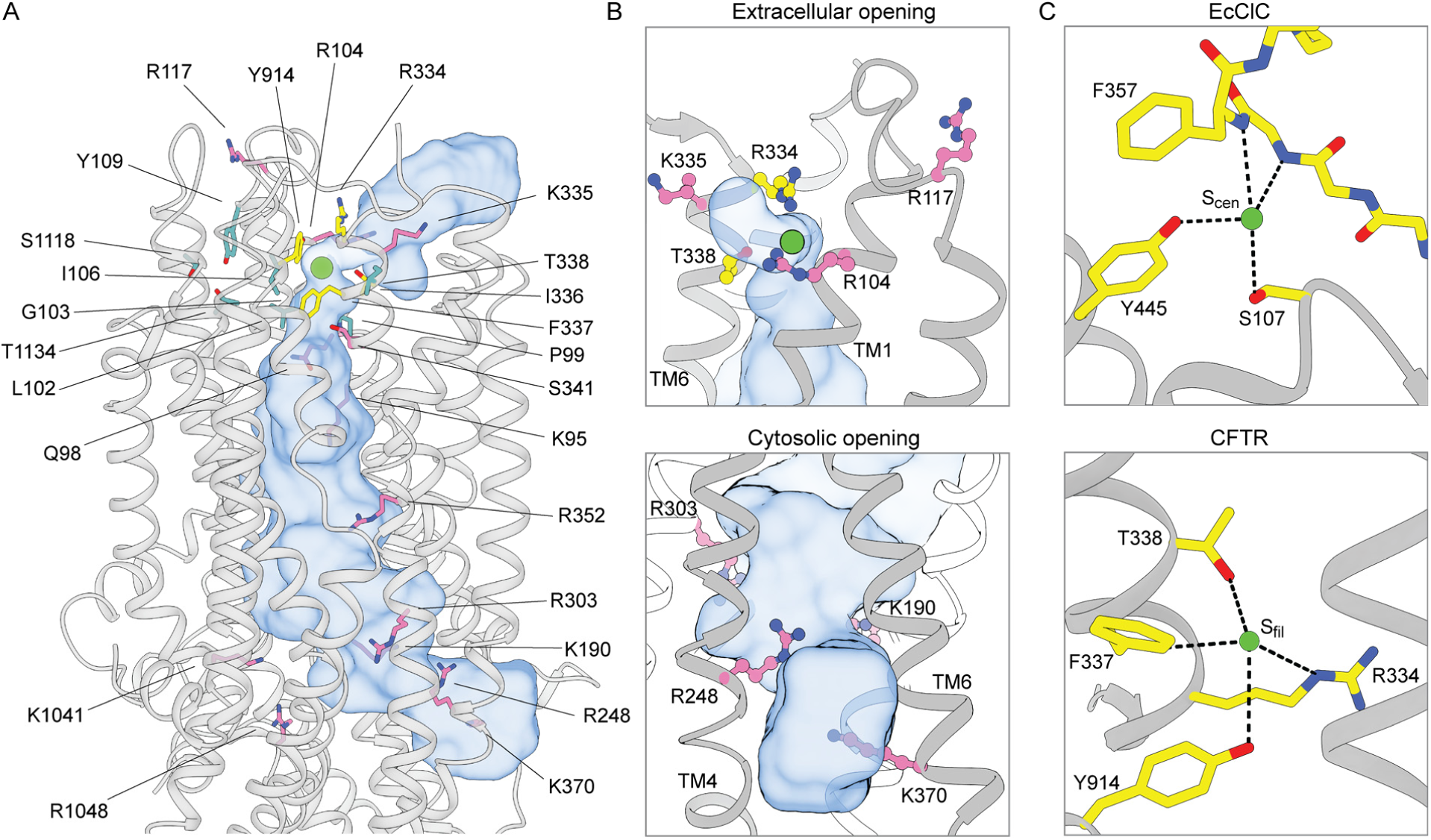
CFTR residues that contribute to ion selectivity and/or ion conduction. (**A**) Residues shown to contribute to ion selectivity or conduction in the literature are mapped onto the three-dimensional structure of CFTR. The pore is shown as a blue surface. Chloride is shown as a green sphere. Residues in the inner and outer vestibules are pink. Residues that directly coordinate chloride at S_filter_ are yellow. Residues of the second coordination sphere of S_filter_ are cyan. (**B**) Closeup views of the extracellular opening (upper panel) and cytosolic opening (lower panel) of CFTR’s pore. The pore is outlined as a blue surface. (**C**) Chloride coordination at the S_cen_ chloride-binding site in wild-type EcClC (upper panel, Protein Data Bank 1OTS) (90) and at the S_filter_ chloride-binding site in CFTR (lower panel).

Previous mutational work has identified a plethora of residues, many are arginine and lysine, that influence CFTR ion selectivity and/or conductance (38, 56, 75–77). Mapping these residues onto the CFTR structure indicates that basic residues, including K95, R104, R117, K190, R248, R303, K335, R352, K370, K1041, and R1048 (Figure 5A, pink), are positioned along the cytosolic and extracellular vestibules, with their side chains exposed to solvent. Different from the residues that directly coordinate chloride at S_filter_ (Figure 5A, yellow), the function of these arginine and lysine residues is to stabilize the partially hydrated anions through electrostatic interactions and to discriminate against cations. The side chains of Q98 and S341 also face the cytosolic vestibule to form anion-dipole interactions with chloride. R334, positioned at the extracellular mouth of the pore, plays a dual role in forming the selectivity filter and attracting anions into the pore through electrostatic interactions. Many other functionally important residues, including P99, L102, I106, Y109, I336, S1118, and T1134 (Figure 5A, cyan), do not directly interact with chloride. Instead, they form a second coordination sphere of S_filter_ that likely contributes to structuring S_filter_ residues with the appropriate geometry to coordinate chloride.

The selectivity filter in CFTR closely resembles that of the CLC chloride channels (78–81), sharing a similar amino acid composition (Figure 5C). In CLCs, the dehydrated chloride at the S_cen_ site is coordinated by positive helix dipoles, main-chain amides, aromatic side chains, and sidechain oxygen atoms of serine and tyrosine residues (78–81). As for T338 and Y914 of CFTR that form similar interactions in S_filter_, substitution of ion-coordinating serine and tyrosine residues in CLC affects ion selectivity, conductance, and open channel block (82, 83). In CLCs, no lysine nor arginine was involved in ion binding. In CFTR, although R334 is part of S_filter_, it interacts with the chloride ion through a weak long-range salt bridge. As discussed earlier by MacKinnon and colleagues (79), a strong Coulomb interaction at a selectivity filter may create a deep energy well and cause a chloride ion to bind too tightly to achieve high conduction rates. Thus, the structures of CFTR and ClCs demonstrate how unrelated ion channels have evolved to select and conduct chloride using common chemical strategies.

It has long been expected that in the fully conductive state, CFTR must exhibit a continuous, open passage across the lipid bilayer, with the narrowest dimensions larger than the permeating ion. Because a probe with radius equal to that of a dehydrated chloride ion cannot fully traverse the extracellular exit between TMs 1 and 6, the conformation of the E1371Q variant was previously interpreted to reflect the rapid flicker closure events that occur within CFTR’s open burst (29, 40, 84, 85). The analysis presented in this work suggests that this notion should be revised. As previously discussed (78), ion channels do not necessarily require a wide pore. Ions can be conducted even when certain parts of the pore on average are marginally narrower than the ions themselves. This is feasible as long as the residues forming the pore possess electrostatic and chemical characteristics that support ion conduction and are flexible to facilitate ion diffusion. In potassium channels, for example, the pore radius between the potassium-binding sites in the selectivity filter is much smaller than that of a potassium ion, yet a conduction rate of 10^8^ per second can be achieved in this configuration (86). Another example is the CLC-1 channel (78), in which the chloride-binding site is separated from the external solvent by a narrow constriction with a 1 Å radius, a configuration similar to that of the external mouth of CFTR. In CFTR, the ion at S_filter_ is separated from the epithelial lumen through a small opening of 1.4 Å, slightly smaller than the 1.7 Å of a dehydrated chloride. Given the high open probability of the E1371Q variant, and the understanding that thermal motions can transiently widen the exit constriction to allow ion passage, we suggest that the structure of the NBD-dimerized E1371Q variant actually reflects the conductive state of CFTR.

## Methods

### Cell culture

Sf9 cells (Gibco, catalogue number 11496015, lot number 1670337) were grown at 27 °C in Sf-900 II SFM medium (Gibco) supplemented with 5% (v/v) heat-inactivated fetal bovine serum (FBS) and 1% (v/v) antibiotic-antimycotic (Gibco). HEK293S GnTI^-^ suspension cells (ATCC CRL-3022, lot number 62430067) were cultured at 37 °C in Freestyle 293 medium (Gibco) supplemented with 2% (v/v) FBS and 1% (v/v) antibiotic-antimycotic. CHO-K1 cells (ATCC CCL-61, lot number 70014310) were cultured at 37 °C in DMEM F-12 medium (ATCC) supplemented with 10% (v/v) FBS and 1% (v/v) GlutaMAX (Gibco).

### Protein expression and purification

CFTR constructs were expressed and purified as previously described (87). Human CFTR with a C-terminal PreScission Protease-cleavable green fluorescent protein (GFP) tag was cloned into the BacMam vector. Recombinant baculovirus was generated using Sf9 cells as previously described (88). HEK293S GnTl^-^ suspension cells, at a density of 2.5 × 10^6^ cells/ml, were infected with 10% (v/v) P3 baculovirus. Protein expression was induced by adding 10 mM sodium butyrate to the culture 12 hours after infection. The cells were cultured at 30 °C for an additional 48 hours and then harvested.

For protein purification, cells were solubilized for 75 minutes at 4 °C in extraction buffer containing 1.25% (w/v) lauryl maltose neopentyl glycol (LMNG), 0.25% (w/v) cholesteryl hemisuccinate (CHS), 200 mM NaCl, 20 mM HEPES (pH 7.2 with NaOH), 2 mM MgCl_2_, 2 mM dithiothreitol (DTT), 20% (v/v) glycerol, 1 μg/ml pepstatin A, 1 μg/ml leupeptin, 1 μg/ml aprotinin, 100 μg/ml soy trypsin inhibitor, 1 mM benzamidine, 1 mM phenylmethylsulfonyl fluoride (PMSF) and 3 µg/ml DNase I.

Lysate was clarified by centrifugation at 75,000*g* for 2 × 20 minutes at 4 °C, and mixed with NHS-activated Sepharose 4 Fast Flow resin (GE Healthcare) conjugated with GFP nanobody, which had been pre-equilibrated in 20 column volumes of extraction buffer. After 1 hour, the resin was packed into a chromatography column, washed with 20 column volumes of wash buffer containing 0.006% (w/v) glyco-diosgenin (GDN), 200 mM NaCl, 20 mM HEPES (pH 7.2 with NaOH) and 2 mM MgCl_2_, and then incubated for 2 hours at 4 °C with 0.35 mg/ml PreScission Protease to cleave off the GFP tag. The eluate was collected by dripping through Glutathione Sepharose 4B resin (Cytiva) to remove PreScission Protease, concentrated, and phosphorylated with protein kinase A (PKA) (NEB) for 1 hour at 20 °C. CFTR was purified by gel filtration chromatography at 4 °C using a Superose 6 10/300 GL column (GE Healthcare), equilibrated with 0.006% (w/v) glyco-diosgenin (GDN), 200 mM NaCl, 20 mM HEPES (pH 7.2 with NaOH) and 2 mM MgCl_2_.

### Proteoliposome reconstitution and planar bilayer recording

The lipids 1,2-dioleoyl-*sn*-glycero-3-phosphoethanolamine, 1-palmitoyl-2-oleyl-*sn*-glycero-3-phosphocholine and 1-palmitoyl-2-oleoyl-*sn*-glycero-3-phospho-L-serine were mixed at a 2:1:1 (w/w/w) ratio and resuspended by sonication in buffer containing 200 mM NaCl, 20 mM HEPES (pH 7.2 with NaOH) and 2 mM MgCl_2_ to a final lipid concentration of 20 mg/ml. 2% (w/v) GDN was added and the mixture was incubated for 1 hour at 25 °C. CFTR was mixed with the lipids at a protein-to-lipid ratio of 1:250 (w/w) and incubated at 4 °C for 2 hours. 14 mg/ml methylated beta-cyclodextrin was added to the mixture. After 4 hours an equivalent amount of methylated beta-cyclodextrin was added to the mixture. This was performed for a total of four additions. Proteoliposomes were pelleted by centrifugation at 150,000*g* for 45 minutes at 4 °C and resuspended in buffer containing 200 mM NaCl, 20 mM HEPES (pH 7.2 with NaOH) and 2 mM MgCl_2_.

Synthetic planar lipid bilayers were made from a lipid mixture containing 1,2-dioleoyl-*sn*-glycero-3-phosphoethanolamine, 1-palmitoyl-2-oleyl-*sn*-glycero-3-phosphocholine and 1-palmitoyl-2-oleoyl-*sn*-glycero-3-phospho-L-serine at a 2:1:1 (w/w/w) ratio. Proteoliposomes containing PKA-phosphorylated CFTR were fused with the bilayers. Currents were recorded at 25 °C in symmetric buffer containing 150 mM NaCl, 2 mM MgCl_2_, 20 mM HEPES (pH 7.2 with NaOH), and 3 mM ATP. Voltage was clamped with an Axopatch 200B amplifier (Molecular Devices). Currents were low-pass filtered at 1 kHz, digitized at 20 kHz with a Digidata 1440A digitizer and recorded using the pCLAMP software suite (Molecular devices). Recordings were further low-pass filtered at 100 Hz. Data were analyzed with Clampfit and OriginPro.

### Patch-clamp electrophysiology

CHO-K1 cells were seeded in 35-mm cell culture dishes (Falcon) 24 hours before transfection. Cells were transiently transfected with BacMam vector encoding C-terminally GFP-fused CFTR, using Lipofectamine 3000 (Invitrogen). 12 hours after transfection, medium was exchanged for DMEM-F12 supplemented with 2% (v/v) FBS and 1% (v/v) GlutaMAX and incubation temperature was reduced to 30 °C. Patch-clamp recording was carried out after an additional 24 hours.

Recordings were carried out using the inside-out patch configuration with local perfusion at the patch. Recording pipettes were pulled from borosilicate glass (outer diameter 1.5 mm, inner diameter 0.86 mm, Sutter) to 1.5–3.0 MΩ resistance. Currents were recorded using an Axopatch 200B amplifier, a Digidata 1550 digitizer and the pClamp software suite (Molecular Devices). Recordings were low-pass-filtered at 1 kHz and digitized at 20 kHz.

For biionic potential measurements, currents were recorded during voltage steps from -80 to 80 mV from a holding potential of 0 mV. Pipette solution contained 150 mM NaCl, 2 mM MgSO_4_, and 10 mM HEPES (pH 7.4 with NaOH). Perfusion solutions contained 150 mM NaX, 2 mM MgSO_4_, and 10 mM HEPES (pH 7.4 with NaOH), where X = F, Cl, Br, I, or SCN. For measurements of SCN block, perfusion solution contained (1-*x_SCN_*)×150 mM NaCl, *x_SCN_*×150 mM NaSCN, 2 mM MgSO_4_, and 10 mM HEPES (pH 7.4 with NaOH), where *x_SCN_* is the mole fraction of monovalent anion that is SCN^-^. MgATP was added where indicated. The ground electrode was connected to the bath via a NaCl-containing agar salt-bridge. Relative permeabilities were calculated from reversal potentials using the Goldman-Hodgkin-Katz equation (69, 70). Reported potentials were corrected for liquid junction potentials. Liquid junction potentials were calculated using the stationary Nernst-Planck equation with LJPcalc (89).

For all measurements, CFTR was activated by exposure to PKA (Sigma-Aldrich) and 3 mM ATP. Experiments were conducted at 25 °C. Displayed recordings were low-pass filtered at 100 Hz. Data were analyzed using Clampfit, GraphPad Prism, and OriginPro.

### Fluorescent size-exclusion chromatography

Adherent HEK293S GnTI^-^ cells were cultured at 37 °C in DMEM-F12 supplemented with 10% (v/v) FBS and 1% (v/v) GlutaMAX in 6-well CellBIND plates (Corning), to 90% confluency. Cells were transiently transfected with BacMam vector encoding C-terminally GFP-fused CFTR, using Lipofectamine 3000 (Invitrogen). 8 hours after transfection, medium was exchanged for DMEM-F12 supplemented with 2% (v/v) FBS and 1% (v/v) GlutaMAX. Cells were cultured for an additional 24 hours at 30 °C, collected by resuspension in Dulbecco’s phosphate buffered saline (Gibco), pelleted by centrifugation, and snap-frozen in liquid nitrogen.

Cells were resuspended in extraction buffer containing 1.25% (w/v) LMNG, 0.25% (w/v) CHS, 200 mM NaCl, 20 mM HEPES (pH 7.2 with NaOH), 2 mM MgCl_2_, 2 mM DTT, 20% (v/v) glycerol, 1 μg/ml pepstatin A, 1 μg/ml leupeptin, 1 μg/ml aprotinin, 100 μg/ml soy trypsin inhibitor, 1 mM benzamidine, 1 mM PMSF and 3 µg/ml Dnase I, and incubated for 1 hour at 4 °C. Clarified lysates were loaded onto a Superose 6 10/300 GL column (GE Healthcare), attached to an analytical high-performance liquid chromatography system with in-line fluorescence detection (Shimadzu). Fluorescence chromatograms were collected at λ_ex_ = 488 nm and λ_em_ = 512 nm.

---

**Figure S1.**
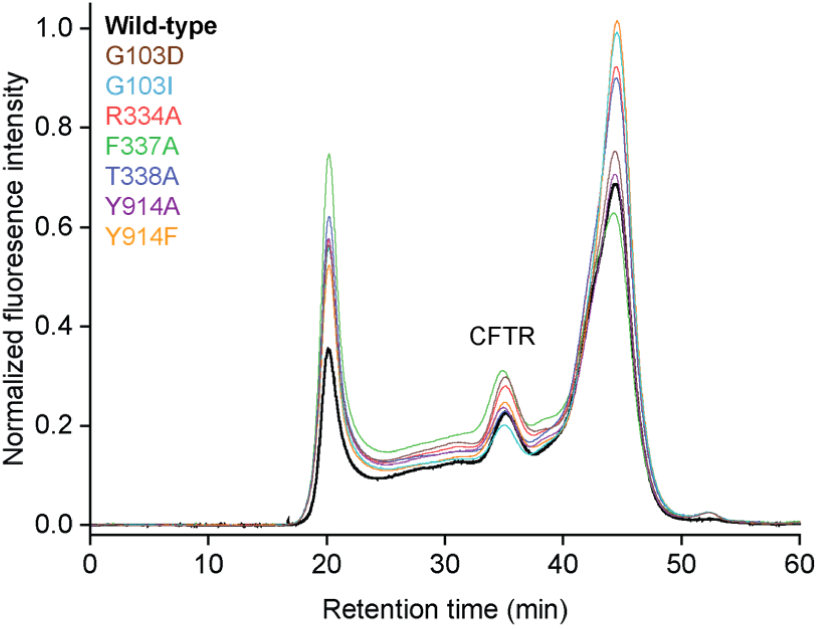
**Fluorescence size-exclusion chromatograms of C-terminally GFP-fused CFTR S_filter_ variants.**

**Figure S2.**
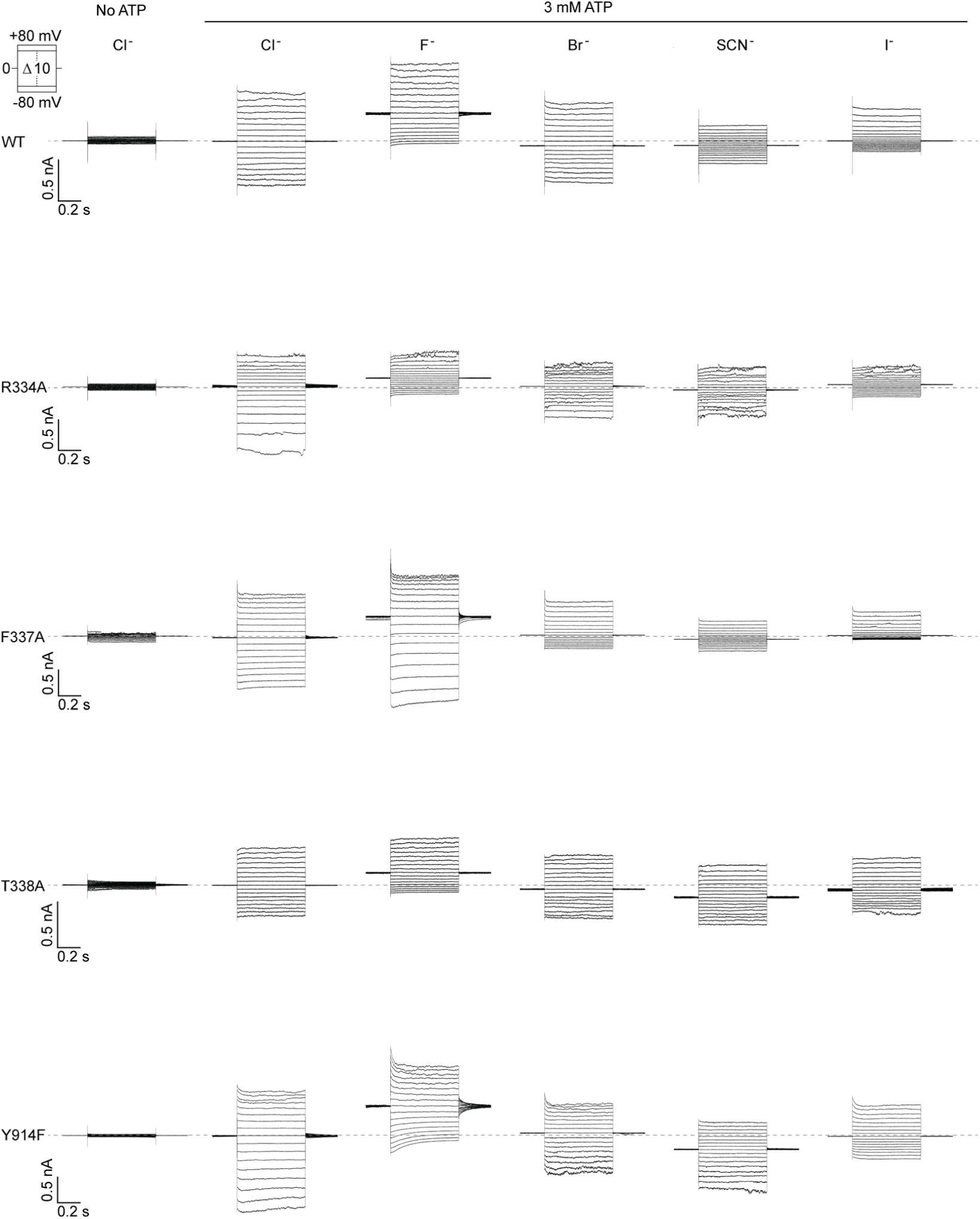
Biionic potential measurements for selectivity filter variants. Example voltage families for PKA-phosphorylated CFTR variants under biionic conditions in inside-out excised patches. The patch pipette contained 150 mM chloride, and the perfusion solution contained 150 mM of the indicated anion. 3 mM ATP was included in the perfusion solution where indicated.

**Figure S3.**
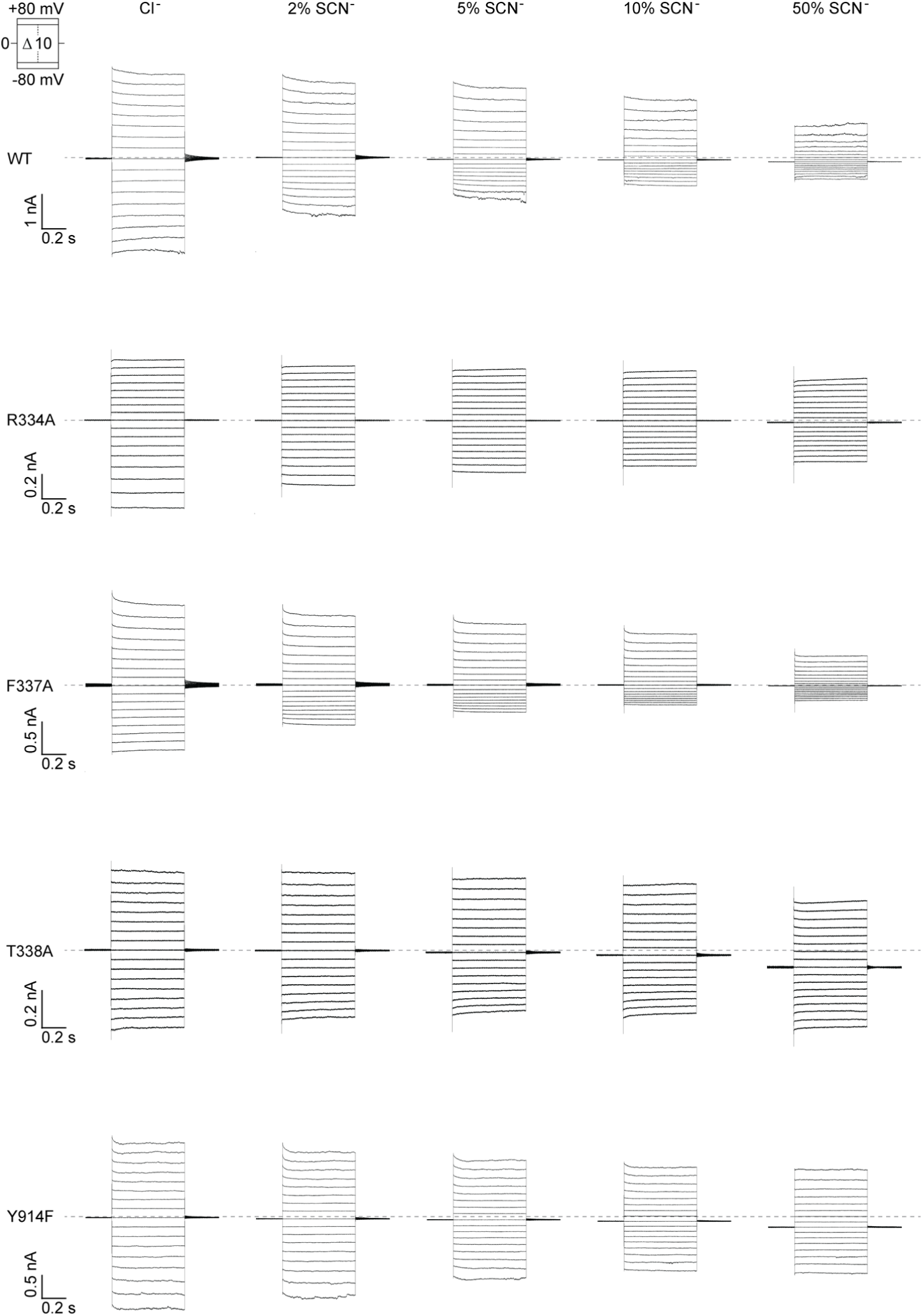
Open channel block of selectivity filter variants by thiocyanate. Example voltage families for PKA-phosphorylated CFTR variants with the indicated mole fractions of thiocyanate. The patch pipette contained 150 mM chloride, and the perfusion solution contained a mixture of chloride and thiocyanate. 3 mM ATP was used in the perfusion solution for all experiments.

## Acknowledgements

We thank members of the Chen and MacKinnon laboratories for helpful discussions, Dr. K. Fiedorczuk for sharing the cryo-EM map before publication, and Dr. L. Csanády for comments on the manuscript. This work was supported by the Howard Hughes Medical Institute (to J.C.).

## Author contributions

J.L. performed the experiments. J.L. and J.C. analyzed the data and wrote the manuscript.

## Data and materials availability

All data and information are available in the main text or the supplementary materials.

